# Ploidy-specific transcriptomes shed light on the heterogeneous identity and metabolism of developing pericarp cells

**DOI:** 10.1101/2023.07.28.550816

**Authors:** Edouard Tourdot, Elie Maza, Anis Djari, Pascal GP Martin, Frédéric Gévaudant, Christian Chevalier, Julien Pirrello, Nathalie Gonzalez

## Abstract

Endoreduplication, during which cells increase their DNA content through successive rounds of full genome replication without cell division, is the major source of endopolyploidy in higher plants. Endoreduplication plays pivotal roles in plant growth and development and is associated with the activation of specific transcriptional programs that are characteristic to each cell type, thereby defining their identity. In plants, endoreduplication is found in numerous organs and cell types and especially in agronomically valuable ones, such as the fleshy fruit (pericarp) of tomato presenting high ploidy levels. We used the tomato pericarp tissue as a model system to explore the transcriptomes associated with endoreduplication progression during fruit growth. We confirmed that expression globally scales with ploidy level and identified sets of genes differentially expressed when comparing ploidy levels at a specific developmental stage. We found that non-endoreduplicated cells are defined by cell division state and cuticle synthesis while endoreduplicated cells are mainly defined by their metabolic activity changing rapidly over time. By combining this dataset with publicly available spatiotemporal pericarp expression data, we proposed a map describing the distribution of ploidy levels within the pericarp. These transcriptome-based predictions were validated by quantifying ploidy levels within the pericarp tissue. This *in situ* ploidy quantification revealed the dynamic progression of endoreduplication and its cell layer specificity during early fruit development. In summary, the study sheds light on the complex relationship between endoreduplication, cell differentiation, and gene expression patterns in the tomato pericarp.

**Significance statement:** The progression of endoreduplication is very dynamic during early fruit development and displays cell layer specific patterns. The integration of ploidy distribution maps with ploidy-specific transcriptome data revealed that gene expression in the pericarp is controlled in a ploidy-specific manner during the early stages of tomato fruit development, resulting in the spatialization of transcriptional programs.

## Introduction

Somatic polyploidy is of common occurrence in various eukaryotic organisms, including insects, mammals, and plants, as an integral part of their normal development (Edgar *et al*., 2014). In higher plants, endoreduplication, which consists in successive rounds of full genome replication without cell division, is the predominant mechanism underlying endopolyploidy (Barow, 2006). Endoreduplication plays critical roles throughout the plant cell life cycle by contributing to diverse developmental processes (Breuer *et al*., 2014; Lee *et al*., 2009) as well as to responses to environmental cues (Lang and Schnittger, 2020). Its functions encompass enabling rapid cell growth (Bhosale *et al*., 2019) and promoting cell specification, as well as contributing to cell and organ morphogenesis (Bramsiepe *et al*., 2010; Roeder *et al*., 2010).

Endoreduplication is associated with the activation of specific transcriptional programs that are pivotal for plant growth and development (Bhosale *et al*., 2018; Pirrello *et al*., 2018). Numerous genes, including regulatory and developmental genes, exhibit expression patterns linked to the endoreduplication state of the cell (Bhosale *et al*., 2018; Pirrello *et al*., 2018; Nagymihály *et al*., 2017). For instance, in the Arabidopsis root cortex, genes involved in cell wall biogenesis display specific expression patterns in 4C cells, suggesting that endoreduplication may contribute to cell wall modifications, and consequently influence cell growth through transcriptional regulation (Bhosale *et al*., 2018; Bhosale *et al*., 2019). In Medicago symbiotic nodule cells, a subset of nodule-specific genes required for symbiotic cell differentiation is expressed in an endoreduplication-dependent manner (Nagymihály *et al*., 2017). Moreover, in the pericarp of fully-grown tomato fruit, beyond a global doubling in gene expression between consecutive ploidy levels, significant changes in the expression of hundreds of genes are also associated with increased ploidy (Pirrello *et al*., 2018). Thus, endoreduplication is essential to establish specific molecular programs that are characteristic to each cell type, thereby defining their identity.

Tomato fruit is composed of the pericarp, which is the fleshy part that derives from the ovary wall, the jelly-like locular tissue surrounding the seeds, and the placenta connecting seeds to the central columella. The size of the fruit is determined by two major processes, cell division and cell expansion, which occur at different rates and durations depending on the tissue type. The pericarp can be further divided into three zones: the exocarp consisting of the three outer layers (E1, E2, E3), the endocarp which corresponds to the two inner layers (I2, I1) and the central layers corresponding to the mesocarp (M and M’) (Renaudin *et al*., 2017). During fruit growth, cell division and expansion activities within the pericarp vary depending on the cell layer (Renaudin *et al*., 2017). Indeed, cell division mainly occurs in the exocarp, although with different rates and orientations depending on the cell layer while cell expansion is more significant in the mesocarp region (Renaudin *et al*., 2017). Concurrent with cell expansion, endoreduplication, which initiates before anthesis, triggers a progressive and important increase in ploidy levels within the pericarp cells. At the end of fruit development, the pericarp consists of a heterogeneous population of cells varying in ploidy levels and sizes. In particular, cells in this tissue exhibit ploidy levels ranging from 2C/4C to 256C and areas ranging from 300 µm^2^ to 45000 µm^2^ (Bourdon *et al*., 2011; Cheniclet *et al*., 2005). Furthermore, endoreduplication in the pericarp is neither randomly nor evenly distributed but rather distributed in a gradient towards the center of the pericarp with the cells within each cell layer having different ploidy levels (Bourdon *et al*., 2011). Pericarp growth is thus probably characterized by specific spatial and dynamic patterns of endoreduplication, which contribute to the proper differentiation of the tissue. Hence, polyploidization probably doesn’t occur uniformly in the population of pericarp cells, resulting in the presence of distinct subpopulations of cells with different DNA content and potentially diverse molecular signatures.

In this work, we used the tomato pericarp tissue and its cells with diverse ploidy levels as a model system to study the transcriptomes associated with the progression of endoreduplication, which is linked to cell differentiation, in actively growing fruit. We obtained transcriptome datasets specific to different ploidy levels from tomato pericarp samples harvested at different timepoints during early fruit development, a transition phase during which fruit growth undergoes more endoreduplication than cell divisions. We confirmed that expression globally scales with ploidy level but also identified hundreds of genes differentially expressed when comparing ploidy levels at a specific developmental stage. These datasets were combined with previously published spatiotemporal pericarp expression data to propose a map describing the distribution of ploidy levels within the pericarp. Subsequently, experimental validation of these transcriptome-based predictions by performing fluorescence *in situ* hybridization to directly assess ploidy levels within the pericarp tissue was performed.

## Results

### Ploidy specific transcriptome during early Tomato fruit development

To investigate the molecular heterogeneity related to endoreduplication during tomato fruit development, we used pericarp from fruits at 6, 9 and 12 days post-anthesis (DPA). These timepoints were chosen because they correspond to an intense period of fruit enlargement, due to a high fruit growth rate (Figure S1), and with significant changes in ploidy levels, in particular the appearance of ploidy levels beyond 8C (16C, 32C and 64C), at the expense of 2C and 4C nuclei (Figure S1). As previously reported for fully grown tomato fruits (Pirrello *et al*., 2018), we used Fluorescence Activated Nuclei Sorting (FANS) to collect nuclei from the pericarp according to their DNA content. At each timepoint, four populations of 100k nuclei, corresponding to the major ploidy classes (>85% of all nuclei), were obtained, namely 2C, 4C, 8C and 16C for 6DPA and 4C, 8C, 16C and 32C for 9 and 12DPA (Figure S1), their RNA was extracted and sequenced by RNA-seq.

Following data quality control and read trimming (Pirrello *et al*., 2018), the reads were mapped to the SLYMIC 1.0 reference genome assembly (BioProject:PRJNA553986) and assigned to annotated genes. Uniquely mapped reads allowed to detect between 13K and 20k transcripts (Table S1). The use of spike-ins controls (ERCC RNA Spike-In Controls) allowed to examine if the transcriptome exhibited a global shift of two-fold change between successive ploidy levels, as reported in fruits at 30DPA (Pirrello *et al*., 2018). A total of 20 Spiked-in control RNAs were detected in at least 28 out of 36 samples (10 in all the samples) and were used to normalize the RNA-seq data by a known, fixed quantity, proportional to the initial number of nuclei. The RNA-seq data normalized using spike-in controls exhibited a clear trend confirming that the transcriptome displays a global 2-fold increase in expression from one ploidy level to the next (Figure S2). Thus, we confirmed that gene expression scales with ploidy level in tomato pericarp, also at early developmental stages of the fruits.

Principal component analysis (PCA) using all expressed genes grouped replicates together and separated the expression profiles between ploidy levels (PC1: 56% of the total variance) and developmental stages (PC2: 24% of the total variance) (Figure S3A). Pairwise comparisons of the expression values were conducted with the DESeq2 package, after standard normalization (Maza, 2016; Maza *et al*., 2013), and fold changes in expression between two ploidy levels at a specific developmental stage were calculated. 18318 genes were identified as non-differentially expressed (non-DE) across all ploidy levels in at least one of the developmental stages, thus having an expression proportional to the ploidy level. Among these, 14843 were non-DE between ploidy levels at all three developmental stages (Table S2). On the other hand, 1938, 1921 and 1878 genes were differentially expressed (DE) in at least one comparison between ploidy levels at 6DPA, 9DPA, and 12DPA, respectively, with an FDR < 1% and log2 FC >1 (Figure S3B and Table S2).

To identify specific expression profiles associated with changes in the ploidy level, the DE genes were clustered via hierarchical clustering, based on the similarity of their expression profiles across ploidy levels. We found that five clusters adequately describing the changes in gene expression during successive rounds of endoreduplication (Figure 1 and Table S3). These clusters exhibited three major expression trends: i) an increased expression with increasing ploidy levels, ii) a decreased expression with increasing ploidy levels, and iii) a peak of expression at specific, intermediate ploidy levels. Specifically, we observed clusters with a peak of expression in 2C (clusters 6-1), 4C (clusters 6-3, 9-1/9-2 and 12-1/12-2), 8C (clusters 9-3 and 12-3), 16C (cluster 6-5) or 32C (clusters 9-5 and 12-5) nuclei. Additionally, we identify a cluster characterized with a high expression in low ploidy levels nuclei, especially 2C and 4C, (cluster 6-2). Conversely, another cluster was found with a high expression in polyploid nuclei, including 8C, 16C and 32C nuclei (clusters 6-4, 9-4 and 12-4).

**Figure 1:**
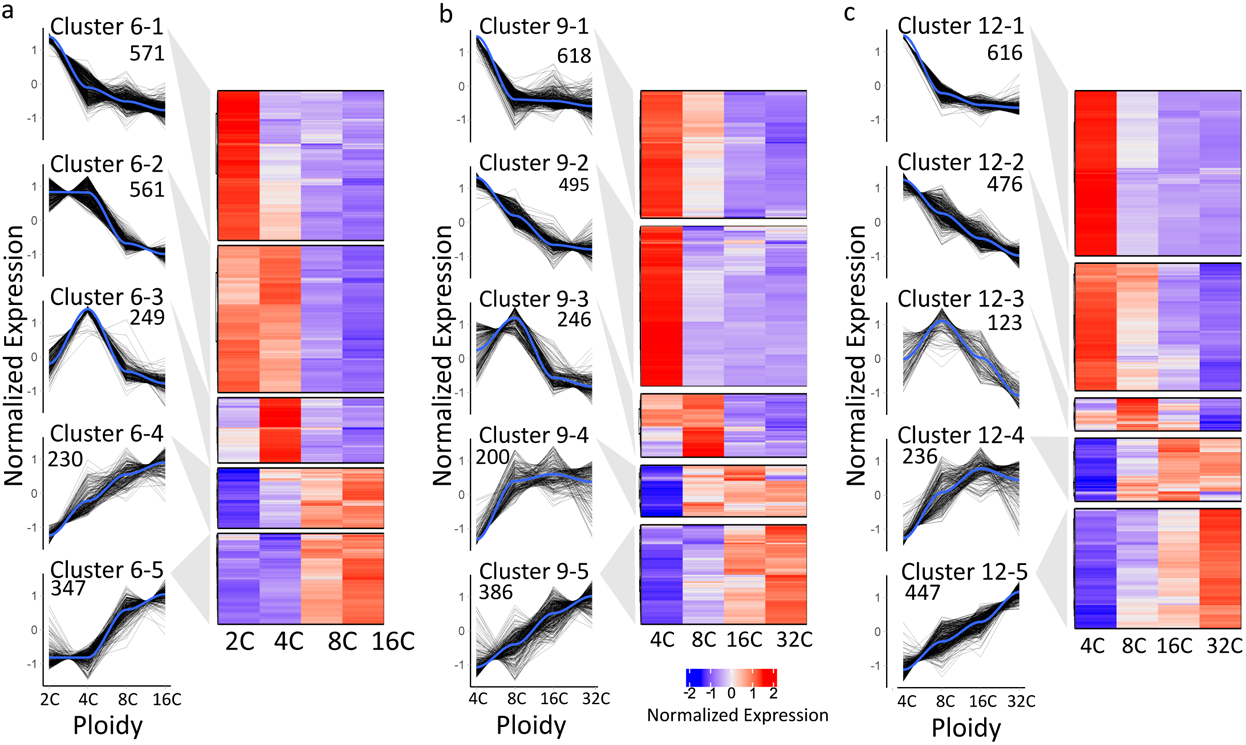
Clustered normalized expression profiles of differentially expressed genes in pericarp of 6, 9 and 12 DPA fruits. Genes were clustered based on their expression patterns in nuclei with four different ploidy levels at 6 (A), 9 (B) and 12 (C) DPA. The graphs and heatmaps present relative gene activities in these nuclei with four different ploidy levels in each gene cluster. Normalized gene expression values (z-scores) are colored according to the color scale at the bottom.

### Non endoreduplicated cells are defined by cell division state and cuticle synthesis

To uncover the biological processes associated with ploidy and developmental stage specific expression profiles, we conducted Gene Ontology (GO) term enrichment analysis in the different clusters. The over-representation of functional categories was assessed using the GO enrichment tool from PLAZA 4.5 (Van Bel *et al*., 2018) (Table S4).

At 6DPA, cluster 6-1 consisted of genes highly expressed in 2C cells while cluster 6-3 comprised genes with high expression specifically in 4C nuclei. These clusters showed enrichment in GO terms mainly related to the canonical cell cycle process (Figure 2). In cluster 6-1, 41 genes were annotated with terms related to “cell cycle”, 91 genes to “DNA metabolic process”, and 55 genes to “Chromatin organization”. In particular, cluster 6-1 included *RBR1* and *E2Fb* genes, encoding important transcriptional regulators of cell division which modulate the expression of numerous genes involved in cell cycle progression (Gutierrez, 2009; Őszi *et al*., 2020). In cluster 6-3, the enriched terms were related to “cell cycle” (29 genes) and “microtubule-based movement” (23 genes), the latter category comprising genes encoding microtubule-associated proteins and kinesin proteins involved in various cellular events such as mitosis and movement of chloroplasts (Li *et al*., 2012). Within the cell cycle term, *CYCLIN* genes such as *CYCA1*, *CYCB2* with roles in the mitotic cycle and the G2/M transition (Menges *et al*., 2005; Sabelli *et al*., 2014), *CYCB3* and *the G2/M specific CYCLIN DEPENDENT KINASE B (CDKB2)* (Menges *et al*., 2005) were identified.

**Figure 2:**
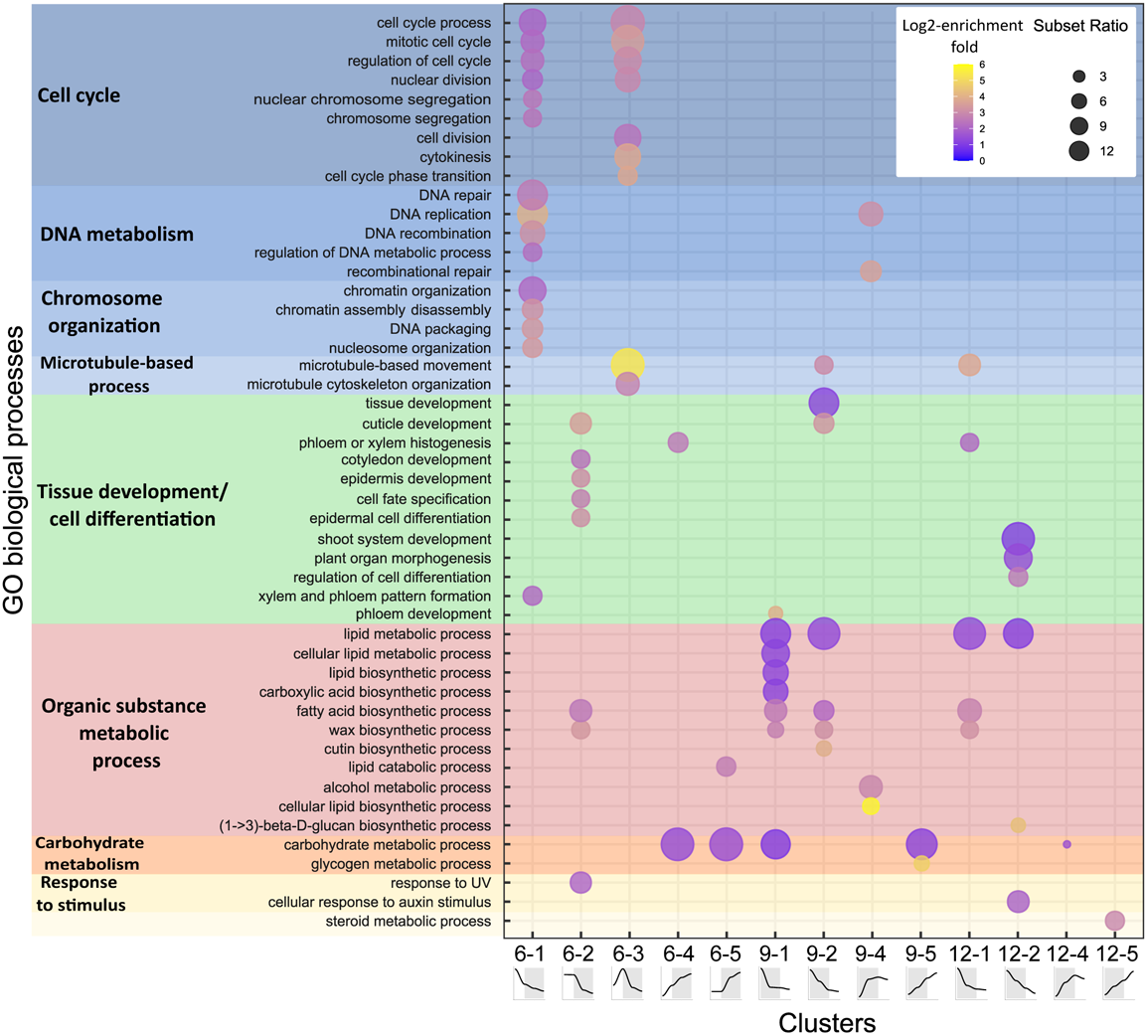
Major functional categories overrepresented in the list of DEGs according to their expression profiles. Gene ontology (GO) enrichment analyses of the DEGs from the 5 clusters per time point. The dot size (subset ratio) is representative of the number of DEGs associated with the process and the fold enrichment is according to a heat map (dot color).

Cluster 6-2 contains genes with highest expression in 2C and 4C nuclei and showed enrichment in terms associated with tissue development (“epidermis development”, 25 genes) and cell differentiation (“epidermal cell differentiation”, 10 genes). Additionally, terms related to the formation of cuticle, an essential hydrophobic extracellular layer protecting the outermost surface of fruits, including “cuticle development” (17 genes), “fatty acid biosynthetic process” (19 genes) and “wax biosynthetic process” (11 genes) (Figure 2) were found in this cluster. Similar enrichment of cuticle related terms was observed at 9 and 12 DPA in clusters 9-1 and 9-2, 12-1 and 12-2 which also showed peak expression in 4C nuclei (Figure 1 and Figure 2).

As in cluster 6-3, the term “microtubule-based movement” was found enriched in clusters 9-1 and 12-1 indicating that this process is particularly relevant in the 4C cells. Cluster 9-1 was enriched specifically in the term “carbohydrate catabolic process” with 14 genes related to glycolysis. In cluster 12-2, terms related to organ morphogenesis were found enriched, involving 50 genes mainly associated to response to hormones, particularly to auxin.

To summarize, non endoreduplicated cells (2C and 4C) at 6 DPA are actively dividing and synthesize the fruit cuticle (clusters 6-1 to 6-3). The genes found in clusters 6-1 and 6-3 suggest that the 2C nuclei could correspond to cells in late G1 phase (G1/S transition), while genes found in 4C nuclei would mainly correspond to cells in late G2 phase, possibly at the G2/M transition. At 9 and 12 DPA, 4C cells appeared to be primarily involved in cuticle synthesis and differentiation processes, with no enrichment of cell cycle related genes indicating a decline in cell division and a likely commitment to endoreduplication for a substantial fraction of the 4C cells.

### Endoreduplicated cells are mainly defined by their metabolic activity changing rapidly over time

Genes mainly expressed in endoreduplicated cells were found in clusters 6-4 and 6-5 at 6DPA, clusters 9-3 to 9-5 at 9DPA and 12-3 to 12-5 at 12DPA (Figure 1).

At 6DPA, genes with highest expression levels in 8C and 16C endoreduplicated nuclei were found in clusters 6-4 and 6-5 (Figure 2). Cluster 6-4 was enriched for genes involved in “carbohydrate metabolic process”, comprising 22 genes involved in central metabolism, particularly related to sucrose metabolism, glycolysis, and the pentose phosphate pathway. In cluster 6-5, genes involved in catabolism and synthesis of complex sugars were enriched, including “carbohydrate metabolic process” (35 genes) and “polysaccharide metabolic process” (16 genes). Among these genes, 10 encoded proteins involved in starch synthesis, secondary cell wall synthesis and modifications. When combining genes from clusters 6-4 and 6-5 for the GO enrichment analysis, the term “transmembrane transport” (41 genes) including sugar and auxin transporters became significantly enriched (Table S4) suggesting an active symplastic movement of molecules in these cells, in parallel with increased carbohydrate metabolism.

At 9 DPA, cluster 9-4 and 9-5 contained genes highly expressed in 8C, 16C and 32C (Figure 2). As in cluster 6-5, one of the two enriched GO terms found in cluster 9-5, “carbohydrate metabolic process” (32 genes), was associated with sugar and starch metabolism, as well as secondary cell wall synthesis. Interestingly, in cluster 9-4, the terms “DNA replication” (9 genes) and “recombinational repair” (6 genes) were enriched with genes encoding MINI CHROMOSOME MAINTENANCE proteins (MCM 2/4/6) and a CELL DIVISION CONTROL PROTEIN 6 HOMOLOG (CDC6). Some MCM proteins have been associated with the endocycle in mammals and maize endosperm (Edgar and Orr-Weaver, 2001; Sabelli *et al*., 2013) while CDC6 proteins have been associated with both the cell cycle and the endocycle replication processes (Castellano *et al*., 2001; Gutierrez, 2009). Although cluster 9-3 contains 243 genes, we found no significant enrichment of GO biological process terms in this cluster, suggesting that the functions of these genes are yet poorly characterized.

Cluster 12-4 was enriched for genes involved in *“carbohydrate metabolic process”* (28 genes), with genes implicated in secondary cell wall synthesis and modifications and sugar metabolism (Figure 2). Cluster 12-5 was only enriched in *“steroid metabolic process”* (9 genes) including three genes implicated in brassinosteroid synthesis or signaling (Figure 2). Finally, cluster 12-3 did not exhibit any significant enrichment of GO biological processes.

Overall, our RNA-seq data suggests that endoreduplicated cells in tomato fruit pericarp at 6 DPA have increased sugar metabolism, likely related to high metabolic activity during the cell proliferation phase, while the increased expression of cell wall-related genes could account for cell growth occurring in the mesocarp (Terao *et al*., 2013; Renaudin *et al*., 2017). At 9DPA and 12 DPA, endoreduplicated cells (8C, 16C and 32C) were already engaged into secondary metabolism pathways probably responsible for colour of tomato fruits in addition to their central metabolism activity. Thus, our data show that the transcriptome of pericarp cell depends on both the ploidy level of the cell and the developmental stage of the fruit.

### Genetic programs conserved between ploidy levels but also shifting to higher ploidy levels over time: diverse cell populations contribute to cuticle synthesis

The analysis of the enriched GO categories revealed that some clusters displaying the same ploidy-dependent changes in expression were enriched in similar GO terms. To evaluate which genes displayed consistent ploidy-dependent changes in expression, we analyzed the overlap between clusters and its statistical significance (Figure 3A). Indeed, significant overlaps were mainly observed on the diagonal of the matrix, indicating that clusters with similar expression trends exhibited consistent, ploidy-dependent gene expression patterns (Figure 3A). This observation indicates that, at 6, 9 and 12 DPA, common genes are consistently regulated as the process of endoreduplication takes place and the ploidy of the cells increases. For each cluster, except 6-1 (high expression in 2C nuclei, not studied at 9 and 12DPA), we found at least one significant intersection among the clusters.

**Figure 3:**
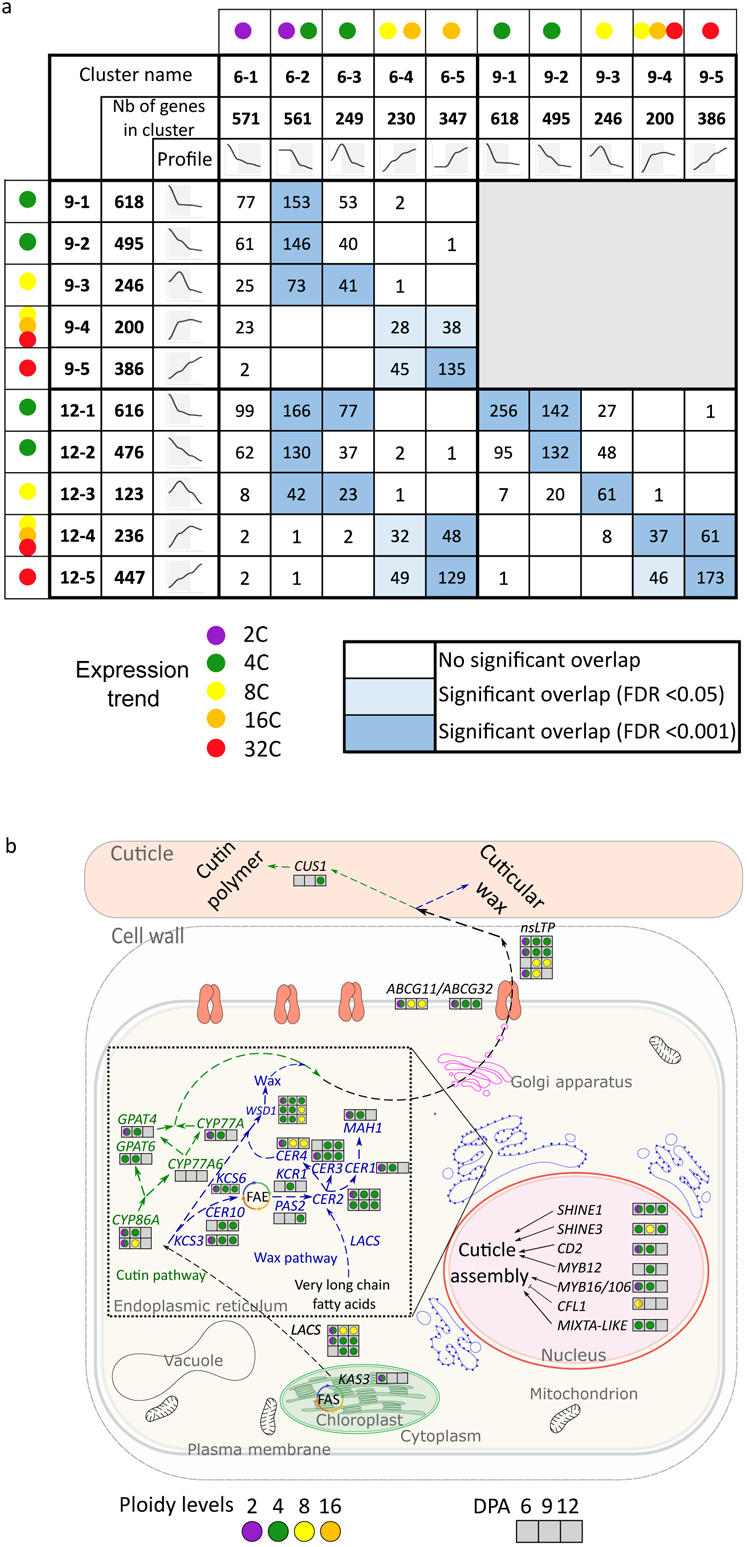
Overlap between clusters of differentially expressed genes in pericarp of 6, 9 and 12 DPA fruits. a. Overlap between genes present in the different clusters at 6DPA, 9DPA and 12DPA. The number of genes in each overlap is shown for each comparison between two clusters. Empty boxes correspond to comparisons with no overlap. Significant overlap for P <0.05 is highlighted by light blue and for P < 0.001 by dark blue. b. Cutin and wax synthesis genes pathways are represented with only transcriptional regulators found differentially expressed in the clusters. Expression peaking at a specific ploidy level is symbolized by a colored circle (or two-colored half circles for expression peaking in two ploidy levels) in a box corresponding to a specific stage (6DPA, 9DPA and 12DPA). Abbreviations: KCS = KETOACYL-CoA SYNTHASE, CYP = CYTOCHROME P450, nsLTP = nonspecific LIPID TRANSPORT PROTEIN, CD2 = CUTIN DEFICIENT 2, MAH = MIDCHAIN ALKANE HYDROXYLASE, KCR = β-ketoacyl reductase, GPAT = GLYCEROL-3-PHOSPHATE ACYLTRANSFERASE, ABCG = ATP Binding Cassette G, CER = ECERIFERUM, WSD = WAX ESTER SYNTHASE, LACS = LONG-CHAIN ACYL-CoA SYNTHETASE, CFL1 = CURLY FLAG LEAF 1, PAS = PASTICCINO, CUS = CUTIN SYNTHASE.

In clusters displaying highest expression in 4C nuclei, GO terms related to cuticle synthesis were systematically enriched. Among the 50 key genes known to be involved in cuticle synthesis, transport, and degradation in tomato fruits, 40 were found in at least one of the clusters and were mainly expressed at high levels in 4C nuclei (Figure 3B and Table S5, (Lashbrooke *et al*., 2015; Petit *et al*., 2017; Trivedi *et al*., 2019)). Notably, 4C nuclei in fruits at 6, 9 and 12DPA, expressed all known genes required for both cutin and wax synthesis such as *SlGPAT4* (cluster 6-2/9-1) and the three tomato *WSD1* orthologs (clusters 6-2/6-3/9-1/9-2/12-1, (Li-Beisson *et al*., 2009; Petit *et al*., 2016)), as well as known transcriptional regulators like SlSHINE1 and SlSHINE3 (Clusters 6-2/9-1/12-1, (Shi *et al*., 2013)) and ABCG’s and nsLTP’s transporters involved in exporting cuticle components to the extracellular space (Fabre *et al*., 2016; Lashbrooke *et al*., 2015). Interestingly, this cuticle synthesis activity was not only limited to 4C cells (Figure 3B), but also occurred in 2C cells at 6 DPA, which are also characterized by a high expression of genes involved in cell cycle activity. At the later 9- and 12 DPA stages, 8C cells were distinguished by their high expression of genes involved in sugar metabolism. This observation strongly suggests a shift in the metabolic activities of low and high polyploid cells with endoreduplication being associated to specific regulations of the transcriptome that result in highly polyploïd cells involved in carbohydrate synthesis and transport.

### Inferring spatial distribution of ploidy levels from transcriptome data

The cuticle is known to be synthesized in the epidermal cell layers of aerial plant organs including leaf and flower petals (Fernández *et al*., 2016). This also applies to tomato pericarp, since the three outer epidermal cell layers are the main sites of cuticle production (Domínguez *et al*., 2008; Segado *et al*., 2016). The expression of genes such as *SlSHINE* 1 and *SlSHINE* 3 which are master activators of most of the cutin and wax synthesis genes, is highest in the epidermal cells (Shi *et al*., 2013; Al-Abdallat *et al*., 2014). In our transcriptome data, most of the genes related to cuticle synthesis were highly expressed in 2C and 4C nuclei at 6 DPA and up to 8C nuclei at 9DPA and 12DPA, with 8C cells specializing in cutin production (Figure 3B and Table S5). This observation indicates that the epidermal cell layers at 6DPA first consist of non endoreduplicated cells but their composition rapidly changes with 8C cells present in the epidermis of 9 and 12DPA fruits. Except for this particular case, no other molecular information from our transcriptome data alone allowed to reallocate the cells with different ploidy within the pericarp tissue. To evaluate how much our transcriptome data is related to the spatial distribution of polyploid cells within the pericarp tissue, we compared our ploidy-dependent transcriptome data to genome wide expression data from pericarp cell type/tissue obtained by Laser Microdissection (LM) from the M82 Tomato cultivar (Shinozaki *et al*., 2018). For this comparison, we used the 2645 so-called “Hub genes” from the LM dataset, which are genes having the strongest correlation within co-expression gene modules identified based on similar expression patterns and associated to specific regions of the pericarp (Shinozaki *et al*., 2018). We used LM expression data from 5 and 10DPA samples, which are the closest from the developmental stages studied in this work (6-9 and 12DPA).

337 genes were found to be common between the DEG from our ploidy-specific transcriptome (3330 genes) and the selected “Hub genes” (2645 genes). We then evaluated the overlap between ploidy-specific clusters and co-expression modules from the LM transcriptome dataset. Significant overlaps were found for 13 ploidy clusters and 9 LM modules (Figure S4). Among the nine modules, five contained genes expressed in specific regions of the pericarp: the vascular tissue for M14, M17 and M19, and the outer epidermis for M20 and M21. The remaining modules contained genes expressed in two regions of the pericarp: genes from M2 were mainly expressed in the outer epidermis and the vascular tissues, while genes from M7 and M8 were expressed in the collenchyma (sub-epidermis layers) and the parenchyma (mesocarp). Genes from M26 were mainly expressed in the collenchyma and to a lesser extent in the vascular tissues.

By combining this spatial information with our ploidy specific expression dataset, we inferred the distribution of ploidy levels within the pericarp (Figure 4 and Figure S4). Genes highly expressed in nuclei with lower ploidy were found in modules expressed in the outer epidermis, in the vascular tissues, and inner epidermis (Figure 4). This spatial distribution of lower ploidy cells (2C and 4C at 6DPA and 4C at 9-12DPA) in the outer epidermis aligns with the GO enrichment analysis that highlighted epidermal related processes such as cuticle synthesis. Clusters of genes with higher expression in high ploidy nuclei significantly overlapped with modules corresponding to genes mainly expressed in collenchyma and parenchyma at 6 and 9DPA. At 12DPA, genes from clusters corresponding to a high expression in highly polyploid cells were found in all pericarp tissues except in the vascular tissue (Figure 4 and Figure S4). This finding indicates that endoreduplication is not restricted to the collenchyma and parenchyma after 12DPA.

**Figure 4:**
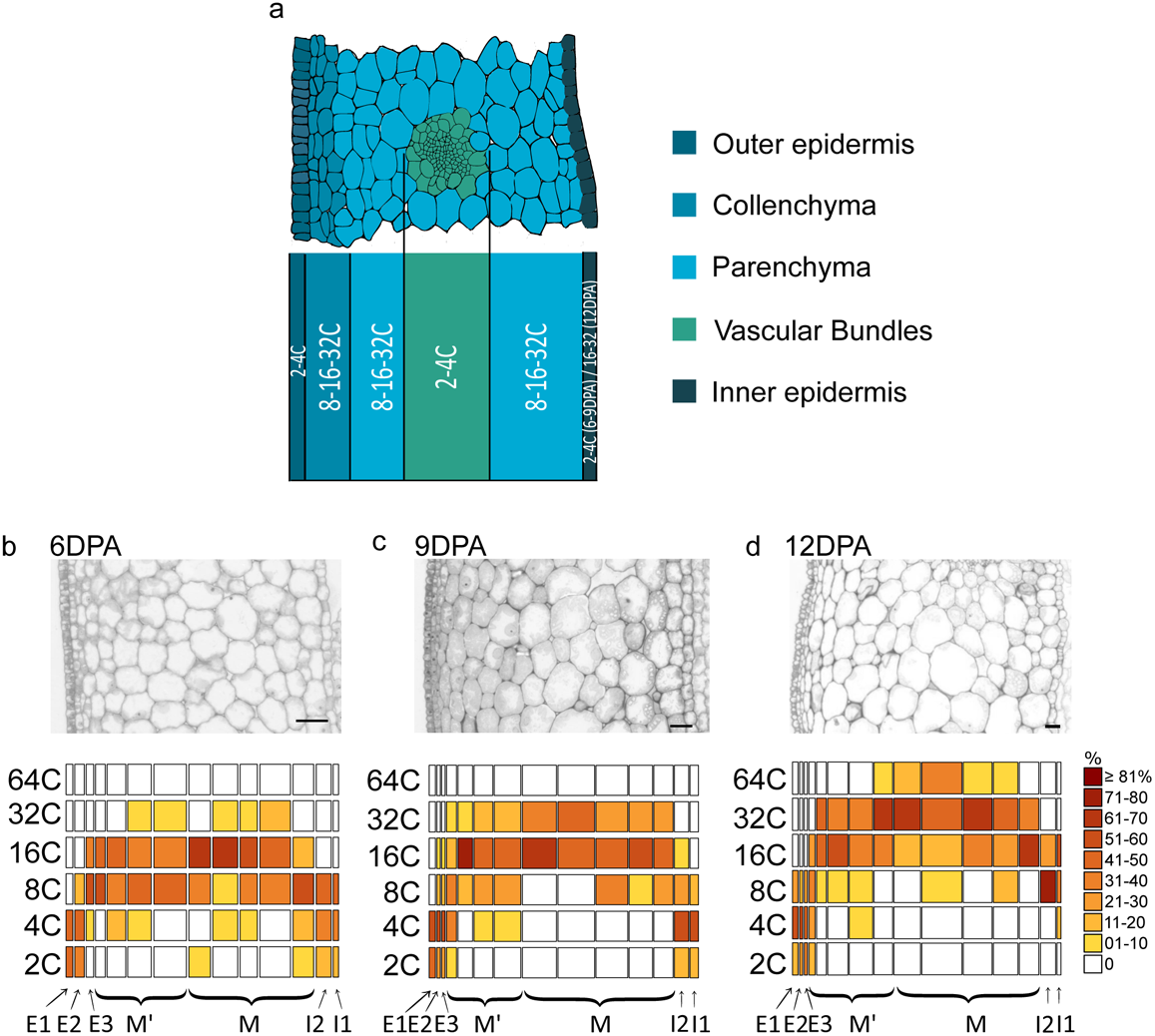
Inferred and measured ploidy distribution in the different cell layers of the fruit pericarp. a. Ploidy map inferred from the comparison between the ploidy-specific transcriptome and the laser microdissected pericarp transcriptome from Shinozaki et al., 2018. b to d. Ploidy maps associated with a representative pericarp section of 6DPA (b), 9DPA (c) and 12DPA (d) fruit. The proportion of each ploidy level per layer is associated with a color scale. n is the number of observed nuclei per layer. E1, E2 and E3 correspond to the outer epidermis (layers 1 to 3). M and M’ layer correspond to distinct mesocarp layers (layers 4 to 12 or 13). I2 and I1 correspond the inner epidermis (layers 13 and 14 or 14 and 15).

### Ploidy distribution patterns during early tomato fruit development

To corroborate these findings, we sought to directly measure the distribution of ploidy levels *in situ* using tomato fruit pericarps at 6, 9 and 12DPA. We employed oligo-Fluorescence *in situ* hybridization (Oligo-F*is*H) to target the 5s rDNA locus, present as a large cluster of tandem repeats on chromosome 1 only (Chang *et al*., 2008). First, we used nuclei with known ploidy levels ranging from 2C to 128C that were sorted by FANS to evaluate the suitability of the 5s rDNA oligo-F*is*H probe for ploidy quantification by counting dot numbers according to the method of Bourdon et *al.* (2011). Clear dot signals were observed in the nuclei incubated with the 5s rDNA FisH probe (Figure S5A) and the number of dots increased with ploidy levels, showing significant differences between two consecutive levels (Figure S5B). Importantly, the counted dots consistently matched the expected ploidy level studied, for example 3.33±0.58 dots and 26±4.21 for 4C and 32C nuclei, respectively. Therefore, the use of the 5s rDNA oligo-F*is*H probe enabled accurate *in situ* quantification of ploidy by classifying the nuclei into unique and specific class of ploidy level based on dot numbers.

Then, ploidy levels were quantified *in situ* in tomato fruit pericarp sections at 6, 9 and 12DPA using the 5s rDNA oligo-F*is*H probe. All hybridized nuclei were imaged, and dot numbers were counted to assign nucleus to a cell layer class according to nomenclature established by Renaudin *et al.,* 2017 (Figure 4B to C). Using this data, we produced ploidy distribution maps for each developmental stage.

At 6DPA, ploidy levels ranged from 2C to 32C (Figure 4B and Figure S6A). 2C and 4C cells were mainly found in the epidermal layers, E1 (∼ 50% of 2C and 4C nuclei), E2 (33% of 2C and 48% of 4C), and in I2 (15% of 2C and 38% of 4C) and I1 (12% of 2C and 38% of 4C). 8C cells were present in all layers except E1, with approximately 50% in E3, M’, I2 and I1, 34% in M layers and 17% in E2 (Figure 4B). 16C cells were absent from the outer and inner epidermal layers but were found from E3 layer to M layers, representing 30% of the cells in E3, 44% in M’ and nearly 50% in most M layers. Among the M layers, the most internal layers, 8, 9 and 10, had the highest proportion of 16C with 62%, 70%, 54%, respectively. Additionally, a small percentage (<10%) of 32C cells were observed in M’ and most of the M layers, while layer 11 contained more than 10% of 32C cells (Figure 4B).

In the 9DPA fruit pericarps, ploidy levels ranging from 2C to 32C were observed, but the distribution and proportion of these cells changed compared to 6 DPA (Figure 4C and Figure S6A). The proportion of 2C, 4C and 8C cells did not undergo drastic changes in the outer and inner epidermal layers compared to 6DPA. However, in the mesocarp layers, the proportion of 8C cells decreased to 29% in M’ and ∼20% in M compared to 6DPA (Figure 4C). As expected, this decrease was accompanied by an increase in the proportion of 16C and 32C nuclei. Specifically, 16C was the most common ploidy level observed in cells from the M’ layers (47%) and the M layers (51% of the cells). Finally, 32C cells were found in all mesocarp layers with proportions of 12% in M’ and 33% in M layers with a maximum of 50% in the layer 9 (Figure 4C).

At 12DPA, ploidy levels ranged from 2C to 64C (Figure 4D and Figure S6A). The outer epidermal cell layers, similar to the 9DPA fruit, were predominantly composed of 2C (20%) and 4C cells (55%), but 8C cells became present in all epidermal layers. In I1, 4C cells accounted for 11% while 2C cells were absent from the I2 and I1 layers (Figure 4C). Most cells in the inner epidermal layers were endoreduplicated, with 8C or 16C nuclei representing 89% in I1 and 100% in I2. 8C cells were mainly present in the outer epidermal layers (21%), in I2 layer (75%) of the cells and in I1 (33%). Progression of endoreduplication was clear in M’ and M layers when comparing 12DPA to 6 or 9DPA pericarp. The proportion of 8C cells decreased, accounting for approximately 10% of the cells (Figure 4D). In the M’ layers, around 40% of the cells were 16C, and the remaining consisted of 32C (approximately 37%), except for the inner M’ layer (layer 8), which had a higher proportion of 32C (61%) and even one 64C nucleus was observed (Figure 4D). In the M layers, cells were predominantly 32C (approximately 57%) and the combined proportions of 32C and 64C reached 70 %. 64C cells accounted for less than 10% of the cells in the M layers, with the highest proportion observed in the inner M layer (layer 10), at 33%.

The endoreduplication index (EI), corresponding to the mean number of endoreduplication cycles per nucleus, and the mean endoreduplication (ME) level corresponding to the sum of each ploidy class weighed by its frequency, exhibited specific patterns according to the cell layers (Figure S6B). Between 6- and 9 DPA, these values remained relatively stable in the outer and inner pericarp, but both EI and ME increased in most of these cells at 12 DPA. In M’ and M layers, EI and ME were consistently higher compared to other cell layers at 6DPA, reaching peak values in cells from the M layer at 12 DPA.

Previous studies have shown that an increase in ploidy levels is generally associated with an increase in both nuclear and cell size, including in the tomato pericarp (Robinson *et al*., 2018; Bourdon *et al*., 2011). From the transcriptomic analysis, genes related to cell wall synthesis and modifications were found in the endoreduplicated nuclei at the three timepoints, suggesting their involvement in cell growth during endoreduplication at 6, 9 and 12 DPA. To verify whether these endoreduplicated cells have initiated cell expansion, we measured cell area and nuclear volume for all cells and nuclei with ploidy levels estimated by F*is*H (Figure S6C). Consistently, the largest cells and nuclei were observed in the cells with high ploidy level from the M layer, which are cells that were already present in the ovary Renaudin et al., 2017, Figure S6C). Interestingly, when comparing cells with similar ploidy levels at different developmental stages, we found that median area and nuclear volume of 2C and 4C cells did not vary between stages. However, for cells ranging from 8C to 32C, these parameters increased over time (Figure S6C). Furthermore, within each ploidy class, cell areas and nuclear volumes were variable with for example, 16C cells being as large as 32C cells at the three timepoints (Figure S6C).

In conclusion, the direct quantification of ploidy within the tissue by Fluorescence *in situ* hybridization validates the proposed ploidy distribution based on transcriptomic data, confirming that non-endoreduplicated cells are mainly located in the outer and inner epidermis, while endoreduplicated cells are predominantly found in the mesocarp. Moreover, our detailed ploidy mapping provides important insights into the spatial and temporal dynamics of endoreduplication within the pericarp tissue at developmental stages when endoreduplication becomes predominant compared to cell division, revealing a rapid increase in ploidy levels, particularly in the internal zones of the mesocarp.

### Integration of ploidy tissue distribution with ploidy specific transcriptome reveals specific spatio-temporal patterns of gene expression

To gain insight into the potential intricate spatio-temporal transcriptional regulation within the pericarp, we integrated the ploidy-specific transcriptome data obtained by RNA-seq with the ploidy mapping of the pericarp obtained by Oligo-F*is*H. Assuming a uniform expression within ploidy populations, gene expression values were weighted according to the proportion of cells of each ploidy level within each cell layer of the pericarp. Subsequently, the weighted expression levels were summed within each layer (Figure 5 and Figure S7).

**Figure 5:**
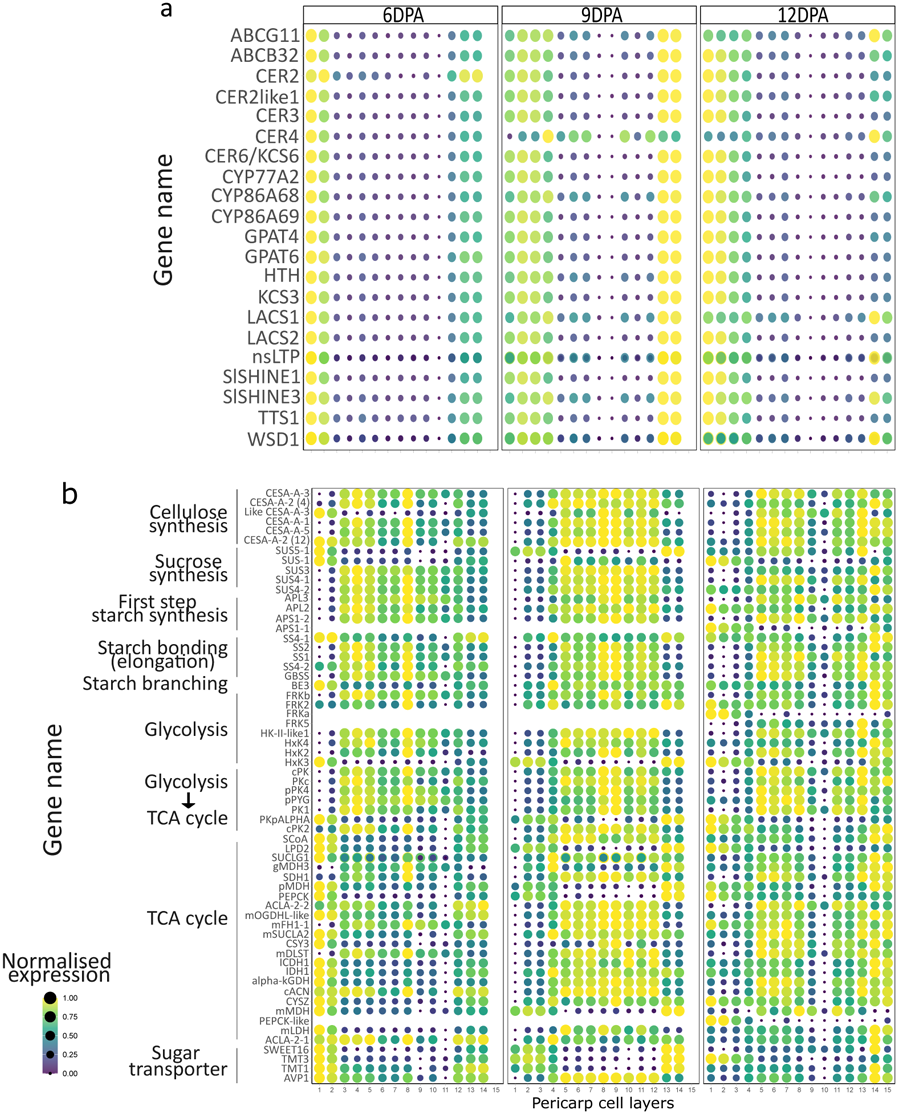
Integration of ploidy transcriptome and ploidy distribution to reveal spatiotemporal ploidy specific gene expression in the pericarp. Spatiotemporal gene expression is obtained from ploidy dependent gene expression weighed by the proportion of each ploidy level in the cell layer. (a) Representation of cuticle synthesis related genes expression in relation with ploidy levels in pericarp cell layer. Abbreviations: ABCG = ATP BINDING CASSETTE G, CER = ECERIFERUM, KCS = KETOACYL-CoA SYNTHASE, CYP = CYTOCHROME P450, GPAT = GLYCEROL-3-PHOSPHATE ACYLTRANSFERASE, HTH: HOTHEAD, LACS = LONG-CHAIN ACYL-CoA SYNTHETASE, nsLTP = nonspecific LIPID TRANSPORT PROTEIN, TTS= TRITERPENOID SYNTHASE, WSD = WAX ESTER SYNTHASE. (b) Representation of metabolism related genes expression in relation with ploidy levels in pericarp cell layer. Both dot size and colors represent normalized expression of genes. Abbreviations: CESA= CELLULOSE SYNTHASE A, SUS= SUCROSE SYNTHASE, APL= AGPase LARGE SUBUNIT, APS= AGPase SMALL SUBUNIT, SS= SOLUBLE STARCH SYNTHASE, GBSS= GRANULE-BOUND STARCH SYNTHASE, BE= STARCH BRANCHING ENZYME, FRK= FRUCTOKINASE, HxK= HEXOKINASE, PK=PYRUVATE KINASE, PYG= GLYCOGEN PHOSPHORYLASE, PKP=PLASTIDIAL PYRUVATE KINASE, SCoA= SUCCINATE--CoA LIGASE, LPD= LIPOAMIDE DEHYDROGENASE, SUCLG= SUCCINYL-COA LIGASE, cMDH= cytosolic MALATE DEHYDROGENASE, PEPCK= PHOSPHOENOLPYRUVATE CARBOXYKINASE, ACLA= ATP-CITRATE LYASE ALPHA-SUBUNIT, OGDHL= OXOGLUTARATE DEHYDROGENASE COMPLEX, FH= FUMARATE HYDRATASE, CSY= CITRATE SYNTHASE, DLST= DIHYDROLIPOAMIDE S-SUCCINYLTRANSFERASE, ICDH=IDH= ISOCITRATE DEHYDROGENASE, alpha-KGDH =Alpha-KETOGLUTARATE DEHYDROGENASE, ACN= ACONITASE, CYS= CITRATE SYNTHASE, MDH= MALATE DEHYDROGENASE, PEPCK= PHOSPHOENOLPYRUVATE CARBOXYKINASE, LDH= LACTATE DEHYDROGENASE, SWEET= SUGARS WILL EVENTUALLY BE EXPORTED TRANSPORTER, TMT= TONOPLAST MONOSACCHARIDE TRANSPORTER, AVP= ARABIDOPSIS VACUOLAR PROTON-PUMPING PYROPHOSPHATASE

Based on these estimated spatial gene expression profiles, genes belonging to clusters 6-1, 6-2 and 6-3, having highest expression in 2C and 4C cells, were found primarily expressed in the outer epidermis, specifically in the layers 1, 2 and 3 (Figure S7). Additionally, genes from clusters 6-2 and 6-3 may exhibit expression in the inner epidermis (layers 13 and 14), while genes from clusters 6-4 and 6-5 are mainly expressed in the mesocarp, spanning layers 4 to 12. Genes from clusters 9-1 and 9-2 are detected in the outer and inner epidermis (Figure S7). In cluster 9-3, gene expression is primarily found in the outer and inner epidermis, with some presence in M’ mesocarp layers, particularly layers 5 and 6, and a single M mesocarp layer (layer 10). Finally, at 9DPA, genes within cluster 9-4 are expressed in cells from M’ and M layers (layers 4 to 12), while cluster 9-5 comprises genes predominantly expressed in M’ layer 5 and M layers. Comparable trends to 9DPA are observed at 12DPA (Figure S7). However, it is noteworthy that at 12DPA, the highest ploidy level assayed in the transcriptome data was 32C, whereas 64C cells are present in five of the 11 mesocarp layers constituting between 3% to 25%. of the cells within the layer Consequently, caution must be taken when interpreting the results for these layers.

This spatial visualization approach was also extended to different sets of genes associated with specific processes expressed at these time points, such as those involved in cuticle synthesis (Figure 5A) or primary metabolism (Figure 5B). This representation highlighted transcriptional disparities among genes within the same family. For instance, 3 *HEXOKINASE* genes were found expressed in the endoreduplicated cells of the mesocarp, while the *HEXOKINASE* 3 gene was expressed in non-endoreduplicated cells, located mainly within the epidermal region. Moreover, changes in the spatial expression over time were observed for gene groups associated with the same pathway, including a substantial portion of genes involved in the TCA cycle. These transcripts initially accumulated in non endoreduplicated cells at 6DPA, but were subsequently expressed predominantly in endoreduplicated cells located in the mesocarp at 9 and 12 DPA.

By integrating the ploidy-specific transcriptome data with the actual distribution of ploidy levels within the pericarp, a comprehensive map of gene expression dynamics throughout the pericarp development can be proposed. This approach enhances our understanding of the complex transcriptional regulation occurring in the pericarp and represents an important first step that will help the interpretation of future single-cell and spatial transcriptomics approaches in this tissue.

## Discussion

Plant cell types and sizes vary from one organ to another, and from one tissue to another, but also within a tissue. The pericarp of tomato, for instance, is made of a heterogenous population of cells that differ not only in size but also in ploidy levels. In this work, by combining FISH for tissue ploidy quantification with FANS-RNAseq based on ploidy levels, we described the evolution of ploidy distribution within the pericarp tissue during early fruit development and defined the transcriptome of polyploid cells. Integration of this data allowed us to propose gene expression maps in this growing tissue.

### The progression of endoreduplication is very dynamic during early fruit development and displays cell layer specific patterns

In a fully grown tomato fruit, the pericarp is composed of cells that vary in size and ploidy level with a gradual increase of both parameters towards the central cell layers (Bourdon *et al*., 2011). Our analysis of ploidy distribution within the pericarp revealed that this gradient is established early during fruit development, as it is already observed at 6 DPA. We also observed mosaic patterns, with most layers containing cells with two to three different ploidy levels, consistent with previous findings obtained in fully grown fruits (Bourdon *et al*., 2011). Our data further revealed that each cell layer undergoes an increase in ploidy levels, albeit with distinct dynamics, resulting in the presence of 8C and 16C cells in the inner epidermal layers and 32C and 64C cells in the mesocarp at 12 DPA. Thus, similar to cell division and expansion processes (Renaudin *et al*., 2017), endoreduplication also exhibits cell layer-specific patterns during early fruit growth. This cell layer specificity in endoreduplication progression may arise from differences in the timing of cells exiting the mitotic cell cycle, similar to what was proposed for the Arabidopsis sepals (Robinson *et al*., 2018) and leaves (Kawade and Tsukaya, 2017), in which cells of various sizes coexist. Kawade (2017) suggested that cells entering into endoreduplication undergo a continuous increase in ploidy, implying that cells entering the endocycle earlier experience a longer period of polyploidization and become larger. Our findings support this hypothesis, since cells in the M layers, which are already larger at anthesis and likely to harbor higher ploidy levels (Renaudin *et al*., 2017), displayed the highest ploidy levels at 12 DPA. Interestingly, while mesocarp cells undergo several rounds of endocycles, exocarp cells mainly undergo cell division but eventually also enter the endocycle. This observation suggests that during early stages of fruit development, a signal may inhibit the onset of endoreduplication (or cell cycle exit) in the external cell layers which is then removed or that a signal activating the endocycle in the central regions, propagates towards the outer and inner regions of the fruit. Therefore, the maps of ploidy distribution reveal that endocycle progression is differentially regulated across the pericarp regions, but not randomly, likely influenced by signals that remain to identify.

### Endoreduplication during early fruit development is associated with cell specialization which is not specific to a ploidy level

Fruit development involves not only cell division and expansion processes essential for growth but also progressive changes in cell composition and metabolism including glycolysis (TCA cycle) during the cell division phase, glycolysis and chloroplastic pathways (starch synthesis and Calvin-Benson cycle) during cell expansion, and enzymatic activities leading to various metabolites accumulating during ripening (Biais *et al*., 2014). Additionally, intense cell wall metabolism is a crucial aspect of fruit growth, supporting both cell division and cell expansion. Analysis of the transcriptome data from nuclei sorted based on ploidy levels revealed that endoreduplicated cells primarily account for these specific metabolic activities, which undergo rapid changes over time. At 6 DPA, endoreduplicated cells appear to be specialized in carbohydrate metabolism related to glycolysis, while at 9 and 12 DPA, endoreduplicated cells (8C, 16C and 32C) are already engaged into more specialized metabolic pathways, in addition to central metabolic activities. In the set of genes associated with carbohydrate metabolism, we found genes involved in both cell wall synthesis and relaxation such as hemicellulose synthesis genes (Rösti *et al*., 2007; Meents *et al*., 2018) and cell wall modification such as Xyloglucan endotransglucosylase/hydrolase (XTH) (Terao *et al*., 2013; Amos and Mohnen, 2019). The expression of these genes in endoreduplicated cells suggests that both the production of new cell walls and the modification of existing cell walls are supporting cell growth. Similar observations have been made for genes related to cell wall synthesis/modification in endoreduplicated cells of Arabidopsis roots (Bhosale *et al*., 2018), indicating that the endocycle may play a significant role in regulating cell wall production and modification to support rapid cell expansion (Bhosale *et al*., 2019).

Interestingly, we found that genes associated with these specific metabolic pathways are expressed in cells with different ploidy levels, implying that both endoreduplication and the position of the cell within the tissue are important for cell specialization in the pericarp.

Overall, while endoreduplication may specify cell differentiation in the pericarp, particularly in relation to carbohydrate metabolism and storage processes (Beauvoit *et al*., 2014), non-endoreduplicated cells are essential for the production of new cells to ensure growth capacity to the pericarp (Liu *et al*., 2003), but also for fruit protection via the synthesis of the cuticle (Hen-Avivi *et al*., 2014). Our data strongly suggests that ploidy level, cell position, and developmental stage within the tissue are critical factors shaping the transcriptome and determining cell fate and differentiation.

### Heterogeneity within the cell populations of similar ploidy: are all 4C or 8C cells the same?

At 6 DPA, 8C cells are found in all cell layers except E1. The transcriptome analysis showed that although numerous genes exhibited a peak of expression in 8C cells, there was no enrichment of specific GO terms, suggesting that the population of 8C cells is heterogeneous, displaying different molecular signatures or lacking specific identity as they transition to higher ploidy levels. When investigating ploidy distribution over time, we found that 4C cells, which constitute the majority of cells in the pericarp at all stages analyzed were present in all four regions of the tissue (E, M’, M and I) at 6DPA. At this stage, the major genes expressed in the 4C nuclei indicate that the cells may be at the G2/M transition, while at 9 and 12 DPA, the transcriptional programs of 4C cells indicate a role in cuticle synthesis and cell differentiation. The 4C cells can undergo different fates: mitosis to produce 2C daughter cells, cell differentiation and entry in a G0 state, or endoreduplication to become an 8C cell. The proportion of each of these types of 4C cells may change over time, and the transcriptional information obtained in our study may represent the larger population at each time point, thus potentially hiding the signals from cells committed to different fates. Flow cytometry is unable to distinguish mitotically dividing G2 cells from endoreduplicating 4C cells or 8C cells with different fates. Therefore, our transcriptomic data likely represents a combination of the expression profiles from different types of cells, with major pathways representing the predominant cell types. To gain further insights and resolve subpopulations with distinct expression programs and roles, it would be valuable to characterize the heterogeneity of these populations using single-cell RNA-seq approach. This would enable the identification of specific cell types and their respective fates. Moreover, integrating spatial transcriptomics with single-cell RNA-seq would be crucial for obtaining a comprehensive spatial representation of gene expression patterns in relation ploidy within the pericarp tissue.

Because ploidy level appears as an important factor that contributes to shape the transcriptome of cells within the pericarp, the data obtained in our study will likely be useful for the interpretation of future datasets obtained by single-cell and spatial transcriptomics approaches.

In conclusion, the integration of ploidy distribution maps with ploidy-specific transcriptome data revealed that gene expression in the pericarp is controlled in a ploidy-specific manner during the early stages of tomato fruit development, resulting in the spatialization of transcriptional programs. The establishment of these specific transcriptiomic profiles warrants further investigation to better understand the underlying mechanisms.

## Experimental procedures

### Plant material and Culture

Tomato plants *Solanum lycopersicum* Mill. Cv. West Virginia 106 (WVA106) were cultivated in greenhouse with photoperiod of 16h light condition at 25°C and 8h night at 20°C and a relative humidity between 70 and 75%. For RNA-seq and F*is*H, individual flowers were tagged on the day of anthesis, fully opened flowers that is wilt the next day. Fruits were harvested at three stages: 6DPA,9DPA and 12DPA.

### RNA preparation from sorted nuclei by FANS

Nuclei were prepared from 20 tomato fruits pericarp at 6DPA and 10 fruits at 9DPA and 12DPA. Nuclei were sorted by FANS as described in (Bourge *et al*., 2018). Pericarp pieces were chopped using a razor blade in 2mL of Gif Nuclear Buffer (GNB: 45 mM MgCl2, 30 mM Sodium-Citrate and 60 mM MOPS pH 7.0, 1% PVP 10.000, 0.1% Triton X-100 and 10 mM sodium metabisulfite (S2O5Na2)) and the suspension was filtered twice through a 48 µm nylon mesh before adding DAPI (4’,6-diamidino-2-phenylindole). 100k nuclei were sorted according to their ploidy levels with a MoFlo ASTRIOS EQ (BECKMAN COULTER) and collected in a tube containing 900µL of TRIzol™ Reagent (THERMOFISHER SCIENTIFIC). Before RNA extraction, 1µL of a dilution 1:1000 of ERCC RNA Spike-In Mix (THERMOFISHER SCIENTIFIC) was added. The mix of nuclei and spike-ins were shortly vortexed and rested for 5mn at room temperature before adding 200µL of Chloroform, agitating the tubes by hand for 15sec and incubating them for 3min at room temperature. After the incubation, tubes were centrifuged for 15mn at 12000g and 4°C, and the upper aqueous phase was transferred (around 840µL) to a fresh tube and mixed with 420µL of 100% ethanol. Then, total RNA was further purified using the RNeasy Micro Kit (QIAGEN) according to the provider’s instructions. A final volume of 14µL of total RNA was obtained.

### RNA sequencing and analysis

Total RNA was used for construction of libraries following the instructions from the Illumina TruSeq Stranded RNA Sample Prep Kit. After library preparation for sequencing, a Duplex-specific nuclease (DSN) treatment was done using the commercial enzyme from EVROGEN following manufacturer’s instructions. Following DSN treatment, libraries were purified with SPRI beads (BECKMAN) and paired-end sequenced (2×150bp) with a HiSeq3000 (ILLUMINA).

Sequence quality controls with FastQC, read trimming with cutadapt (https://doi.org/10.14806/ej.17.1.200) and read mapping with STAR (Dobin *et al*., 2013) was performed as described in Pirrello et al., 2018. We used the SLYMIC 1.0 reference genome along with ERCC sequences for read mapping with STAR (Dobin *et al*., 2013). After mapping, paired reads were assigned to genes or ERCC spike-ins using featurecounts from subread package (Liao *et al*., 2014) and SLYMIC 1.0 annotation. ERCC spike-ins were initially used to normalize gene expression and verify that the global transcriptional activity follows the ploidy level. Differential gene expression analyses were performed with the DESeq2 package (Love *et al*., 2014), after standard normalization, in R/Bioconductor as previously described (Pirrello *et al*., 2018). We considered as differentially expressed the genes that displayed an absolute log2 fold-change > 1 and an FDR-adjusted p-value <0.01. Then DE genes were clustered by hierarchical clustering using z-scores ((expression value) – (mean expression of gene) / (standard deviation of expression of gene)). Distances between genes were calculated using the Euclidean distance, and Ward method applied to square values was used as agglomeration method. Gene ontology enrichment was done using Plaza 4.5 (Van Bel *et al*., 2018) using Bonferroni-corrected pvalue <0.05 and default parameters.

### FISH for ploidy quantification

Nuclei sorted by FANS were recovered on a microscope glass slides (Superfrost) prepared with a 10 µl cushion made of 2M sucrose, 50% GNB and 1% paraformaldehyde. Fruits were sectioned transversally around the equator region and pericarp sections of 100 to 150 µm thickness were obtained using a vibrating blade microtome Microm HM 650V (Microm MicroTech France, http://mm-france.fr/). Two to three slices were placed in a well of a 24 well plate containing a fixative solution (Formaldehyde 4%, DMSO 10% in PBS1X), infiltrated for 5min under vacuum at 20kDA. The FisH protocol was adapted from (Bey *et al*., 2018). Samples were incubated in successive solvent baths for 5min each (2x methanol 100%, 2x ethanol 100%). In the last ethanol baths samples were incubated for at least 45mn at 4°C and then transferred in a solution of 50%ethanol/50%xylene for 30mn at RT. Before proceeding to the next steps, extra care was taken to remove all traces of xylene solution before successive solvent wash baths for 5min at RT each (2x ethanol). Then samples were rehydrated with successive wash baths in water-based buffer for 5min at RT in each bath (2x PBS1X+0.1% v/v Tween20 and 2x SSC2X) and a last incubation was performed in SSC2X in the presence of 100µg/mL of RNAseA for 1h at 37°C, then an equal volume of Hybridization buffer (HB, 50 mM Sodium phosphate pH 7.0, 2x SSC, 50% deionized formamide) was added before incubating the samples for 30mn at RT. The samples were then transfered to a 0.2mL PCR microtube containing 100µL of HB, incubated for 30min before adding 100µL of hybridization buffer with the TexasRed-labelled oligonucleotide (0.5 to 3µL (100µM) of labelled oligonucleotide). Tubes were then incubated in a thermal cycler for 1h at 37°C, then 4min at 88°C followed by a ramp down to 37°C in 3min and incubation at 37°C for 12-15h. The next day, the samples were washed in a stringent bath of HB for 30mn at 42°C followed by 5 washes for 10min in PBS1X, at RT, then 15min in PBS1X+ DAPI (20µg/mL) at RT and 3x 5min at RT in PBS1X. Finally, the samples are transferred to superfrost slides in VECTASHIELD containing DAPI (VECTOR LABORATORIES) under stereomicroscope to assure the flatness of the samples. Finally, the samples were covered with a coverslip (20×20, #1.5) and sealed with nail hardener. The samples were either imaged immediately or after storage for up to several weeks at 4°C with no noticeable change in the signal recovered.

FisH image acquisitions were done with a confocal microscope Zeiss LSM880 with airyscan piloted with Zen Black 2.1 using a x63 objective with oil immersion in airyscan mode. Airyscan raw images were processed using ZenBlue 2.3 with automatic parameters. Then images were analyzed using Fiji (Schindelin *et al*., 2012). After filtering steps corresponding to both gaussian and median filters, nuclei were segmented and the voxel contained in the segmented nucleus were quantified to measure the volume. Cell areas have been determined manually by surrounding cells on their largest diameter. Ploidy counting was also done manually checking in 3D successions of image if close spots correspond to a unique or are duplicated spots.

## Acknowledgments

This work was carried out with the financial support of the Ministère de l’Enseignement Supérieur et de la Recherche (PhD grant to E. Tourdot). We would like to thank Isabelle Atienza-Babin and Aurélie Honoré for taking care of the plants in the greenhouses. The transcriptome sequencing was performed in the GeT-PlaGe facility from GenoToul (https://www.genotoul.fr/).

## Short legends for Supporting Information

**Figure S1:** Tomato fruit diameter-and pericarp ploidy levels during development

**Figure S2:** Global shift of expression in tomato pericarp cells with different ploidy levels

**Figure S3:** Differentially expressed genes between different ploidy levels at 6, 9 and 12 DPA in tomato pericarp cells

**Figure S4:** Overlap between ploidy specific transcriptome and LM pericarp transcriptome of Shirozaki et al., 2018

**Figure S5:** Ploidy quantification on sorted nuclei based on their ploidy using a 5S rDNA targetting oligo-FiSH probe

**Figure S6:** Evolution of ploidy, endoreduplication index and mean ploidy, cell size and nuclear volume in the different pericarp zones over time

**Figure S7:** Integration of ploidy transcriptome and FisH ploidy quantification

**Table S1:** Number of transcripts detected in each sample

**Table S2:** List of differentially expressed genes between different ploidy levels at 6, 9 and 12 DPA

**Table S3:** List of differentially expressed genes in the different clusters

**Table S4:** Overrepresentation of functional categories in the list differentially expressed genes from each cluster at 6, 9 and 12 DPA

**Table S5:** Supplemental table 5: List of key genes described as involved in cuticle synthesis, transport, and degradation in Tomato fruits and their presence in the different clusters of DEG.

## Extended legends for Supporting Information

**Figure S1:** Tomato fruit diameter-and pericarp ploidy levels during development

**Figure S2:** Global shift of expression in tomato pericarp cells with different ploidy levels

(a) MA plots representing the log2 of fold change of reads counts in function of the mean number of normalized read counts between the compared ploidy levels. The red line is indicating the absence of differential expression (Fold change=0). Most genes are expected to have a fold change value around this line if there is no shift of expression between ploidy levels: grey dots. The green dotted line is indicating the expected fold change of expression if there is a shift of expression following ploidy levels. The red dots represent genes having a significant expression fold change. The blue dots corresponds to the spike-ins control.

(b) Fold changes of spike-ins control expression between two ploidies at 6DPA, 9DPA and 12DPA represented as the 95% confidence interval of the fold changes after ANOVA. The dotted lines are indicating the expected fold change between ploidy levels in the case of doubling of expression between two successive ploidy levels.

(a) Principal component analysis (PCA) of ploidy specific transcriptome data. The two first components explain 80% of data variability. PC1, explaining 56% of the total variance, separates the sample according to their ploidy level and PC2, explaining 24% of the total variance, separates the samples according to the developmental stage. Samples are indicated as the corresponding developmental stage (e.g. the three D09.04C correspond to 4C nuclei of 9DPA pericarp). Font colors indicate the ploidy levels: yellow for 2C, green for 4C, orange for 8C, red for 16C and pink for 32C. Grey contours indicate similar developmental stages and blueoverlay similar ploidy levels.

(b) Number of genes up-(yellow) and downregulated (blue) in pairwise comparisons of the expression values (FDR≤0.01, log2FC >1).

Overlap between genes from in the different clusters at 6DPA, 9DPA and 12DPA and Laser Microdissection Pericarp (LM) transcriptome. The number of genes in each overlap is shown for each comparison between two clusters. Empty boxes correspond to comparisons with no overlap.

(a) Representative illustration of nuclear morphology of different ploidy levels. The green dots correspond to the hybridization signal of the 5sr DNA oligo FisH probe. Scale bar = 5µm.

(b) Boxplot representing the counted dots in nuclei sorted based on their ploidy level. The associated significance codes are: 0 ‘****’ 0.001 ‘***’ 0.01 ‘**’ 0.05 ‘

**Figure S7:** Integration of ploidy transcriptome and FisH ploidy quantification

Spatiotemporal gene expression in relation with the ploidy levels in each layer for the five clusters at 6DPA, 9DPA and 12DPA. Spatiotemporal gene expression is obtained from ploidy dependent gene expression weighed by the proportion of each ploidy level in the cell layer for each cluster of genes8 and each stage.

