## Supplementary figures and images for "Ploidy-specific transcriptomes shed light on the heterogeneous identity and metabolism of developing pericarp cells"

### Figure S1

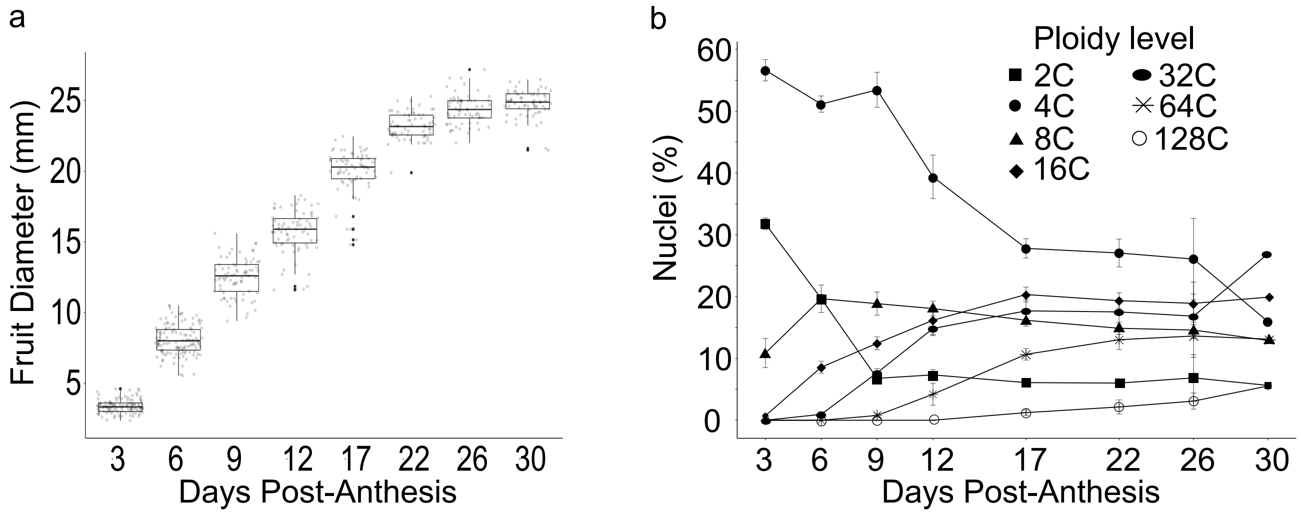

### Figure S2

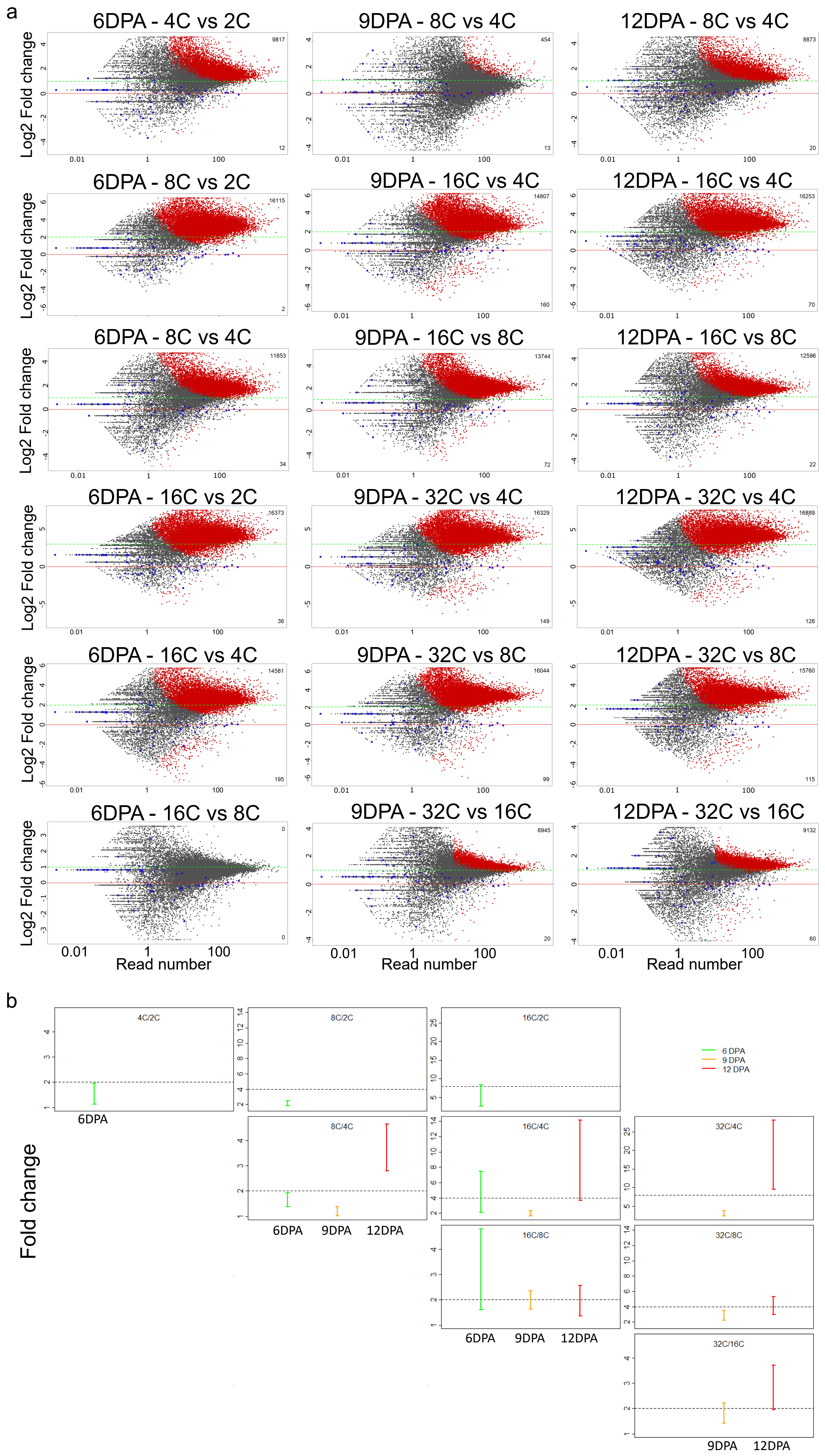

### Figure S3

a

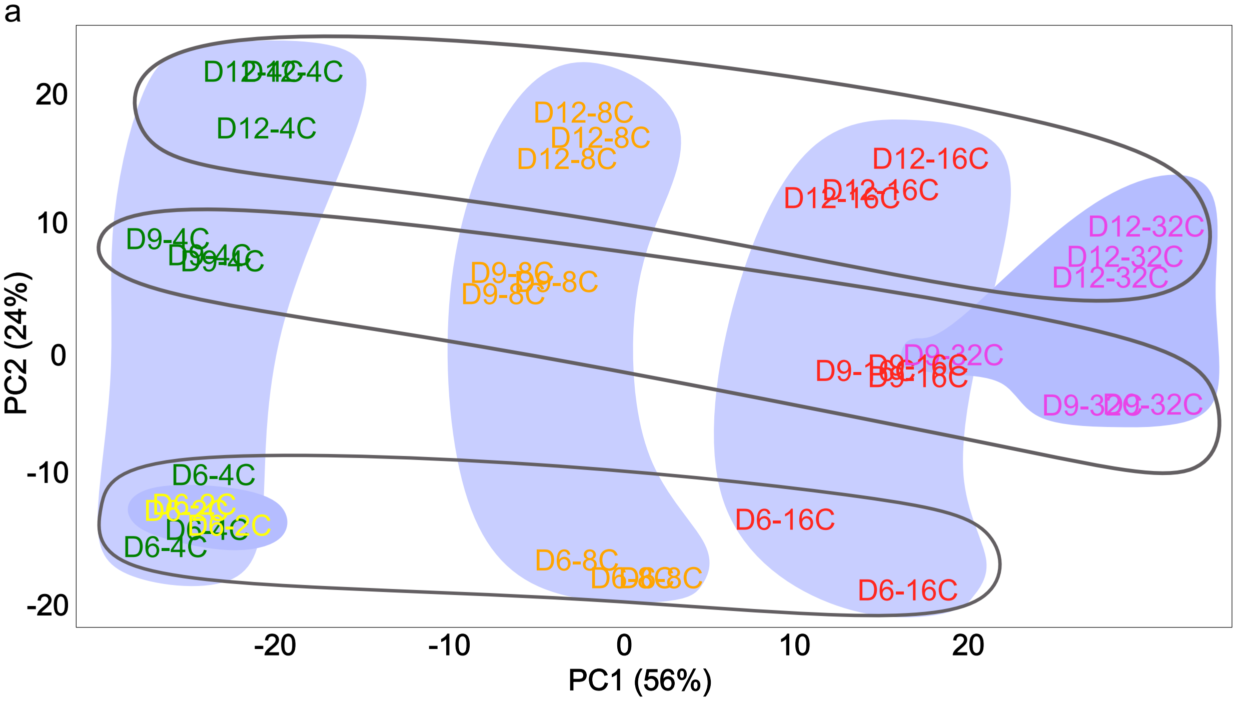


b

| 6DPA (1938 genes) | | | | |
|---|---|---|---|---|
| | **2C** | **4C** | **8C** | **16C** |
| **2C** | | 34 | 297 | 454 |
| **4C** | 96 | | 180 | 300 |
| **8C** | 536 | 383 | | 0 |
| **16C** | 1050 | 831 | 71 | |

| 9DPA (1921 genes) | | | | |
|---|---|---|---|---|
| | **4C** | **8C** | **16C** | **32C** |
| **4C** | | 135 | 371 | 482 |
| **8C** | 220 | | 25 | 82 |
| **16C** | 761 | 222 | | 0 |
| **32C** | 1095 | 493 | 53 | |

| 12DPA (1878 genes) | | | | |
|---|---|---|---|---|
| | **4C** | **8C** | **16C** | **32C** |
| **4C** | | 174 | 401 | 604 |
| **8C** | 185 | | 7 | 78 |
| **16C** | 662 | 113 | | 3 |
| **32C** | 1090 | 473 | 135 | |

### Figure S4

| | | | 6-1 | 6-2 | 6-3 | 6-4 | 6-5 | 9-1 | 9-2 | 9-3 | 9-4 | 9-5 | 12-1 | 12-2 | 12-3 | 12-4 | 12-5 |
|---|---|---|---|---|---|---|---|---|---|---|---|---|---|---|---|---|---|
| | | | 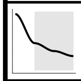 | 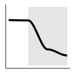 | 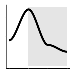 | 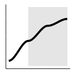 | 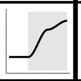 | 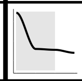 | 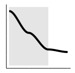 | 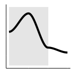 | 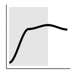 | 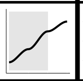 | 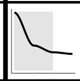 | 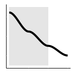 | 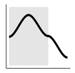 | 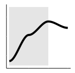 | 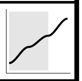 |
| | | | 553 | 542 | 242 | 227 | 358 | 618 | 460 | 241 | 205 | 376 | 606 | 449 | 128 | 246 | 432 |
| **M2** | 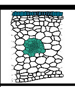 | 156 | 24 | 10 | 31 | | | 19 | 11 | 2 | 4 | 1 | 22 | 5 | 1 | 1 | 1 |
| **M7** | 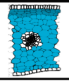 | 41 | | | 1 | 2 | 8 | | | | 2 | 6 | | | | 1 | 8 |
| **M8** | 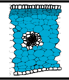 | 66 | | | | 9 | 5 | | | | 4 | 5 | | | | 6 | 8 |
| **M14** | 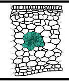 | 29 | 2 | 4 | 1 | | | 3 | 3 | | | | 6 | 3 | | | |
| **M17** | 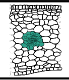 | 36 | 5 | 8 | | | | 6 | 2 | 1 | | | 12 | 6 | | | |
| **M19** | 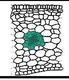 | 42 | 4 | 2 | 2 | | | 11 | 1 | 2 | | | 9 | 8 | | | |
| **M20** | 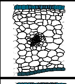 | 49 | 1 | 23 | 9 | | | 28 | 13 | | | | 39 | | | | |
| **M21** | 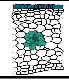 | 17 | 2 | 4 | 1 | | | 6 | 4 | | | | 3 | 5 | | | |
| **M26** | 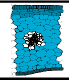 | 36 | | | | 1 | 1 | | | | 1 | 2 | | | | | 5 |

| | |
|---|---|
| | No Significant overlap |
| | Significant overlap (FDR < 0.05) |
| | Significant overlap (FDR < 0.001) |

### Figure S5

a

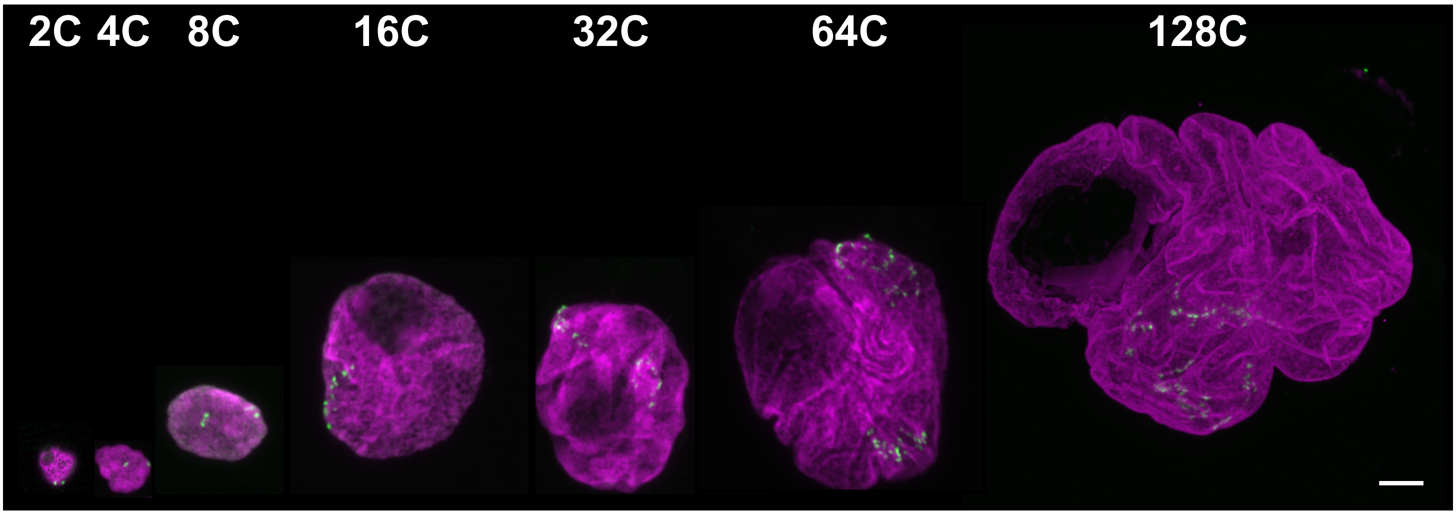


b

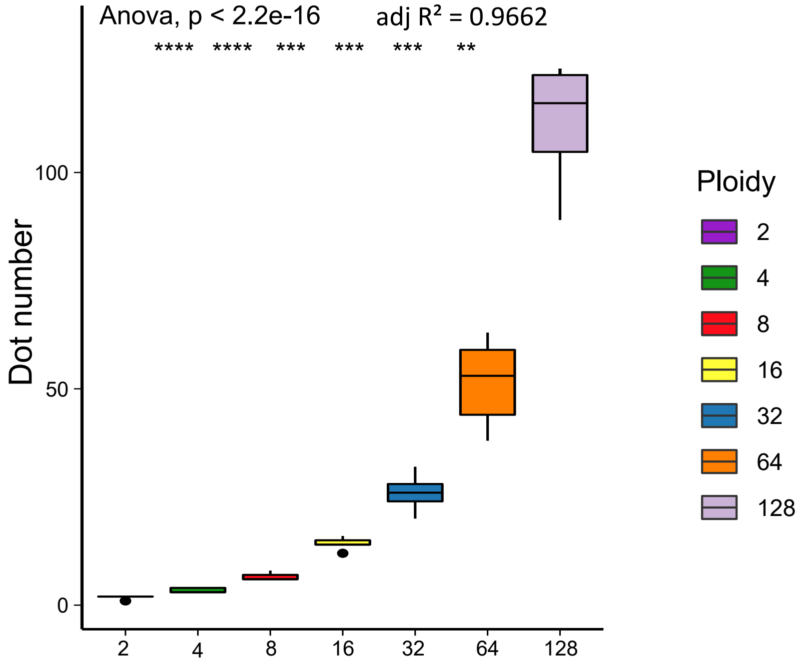


| | | | | | | | |
|---|---|---|---|---|---|---|---|
| 2 | 3,33 | 6,54 | 14,38 | 26 | 51,69 | 111,25 | Average |
| 0,53 | 0,58 | 0,66 | 1,19 | 4,21 | 9,16 | 16,07 | SD |

Ploidy

### Figure S6

a

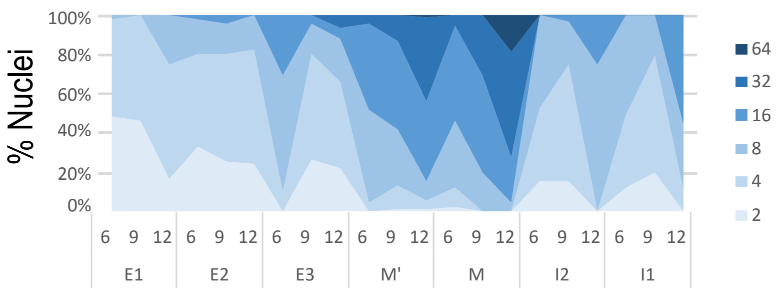


b

| Layer | DPA | EI | mean ploidy |
|---|---|---|---|
| **E1** | 6 | 0,53 | 3,1 |
| | 9 | 0,53 | 3,06 |
| | 12 | 1,08 | 4,66 |
| **E2** | 6 | 0,88 | 4,26 |
| | 9 | 0,98 | 4,6 |
| | 12 | 0,92 | 4,18 |
| **E3** | 6 | 2,2 | 10 |
| | 9 | 0,96 | 4,54 |
| | 12 | 1,3 | 6,84 |
| **M'** | 6 | 2,47 | 12,28 |
| | 9 | 2,57 | 14,18 |
| | 12 | 3,19 | 21,58 |
| **M** | 6 | 2,41 | 12,42 |
| | 9 | 3,07 | 19,04 |
| | 12 | 3,85 | 32,88 |
| I2 | 6 | 1,3 | 5,5 |
| | 9 | 1,12 | 4,92 |
| | 12 | 2,25 | 10 |
| I1 | 6 | 1,38 | 5,76 |
| | 9 | 1 | 4,4 |
| | 12 | 2,45 | 12,04 |

c

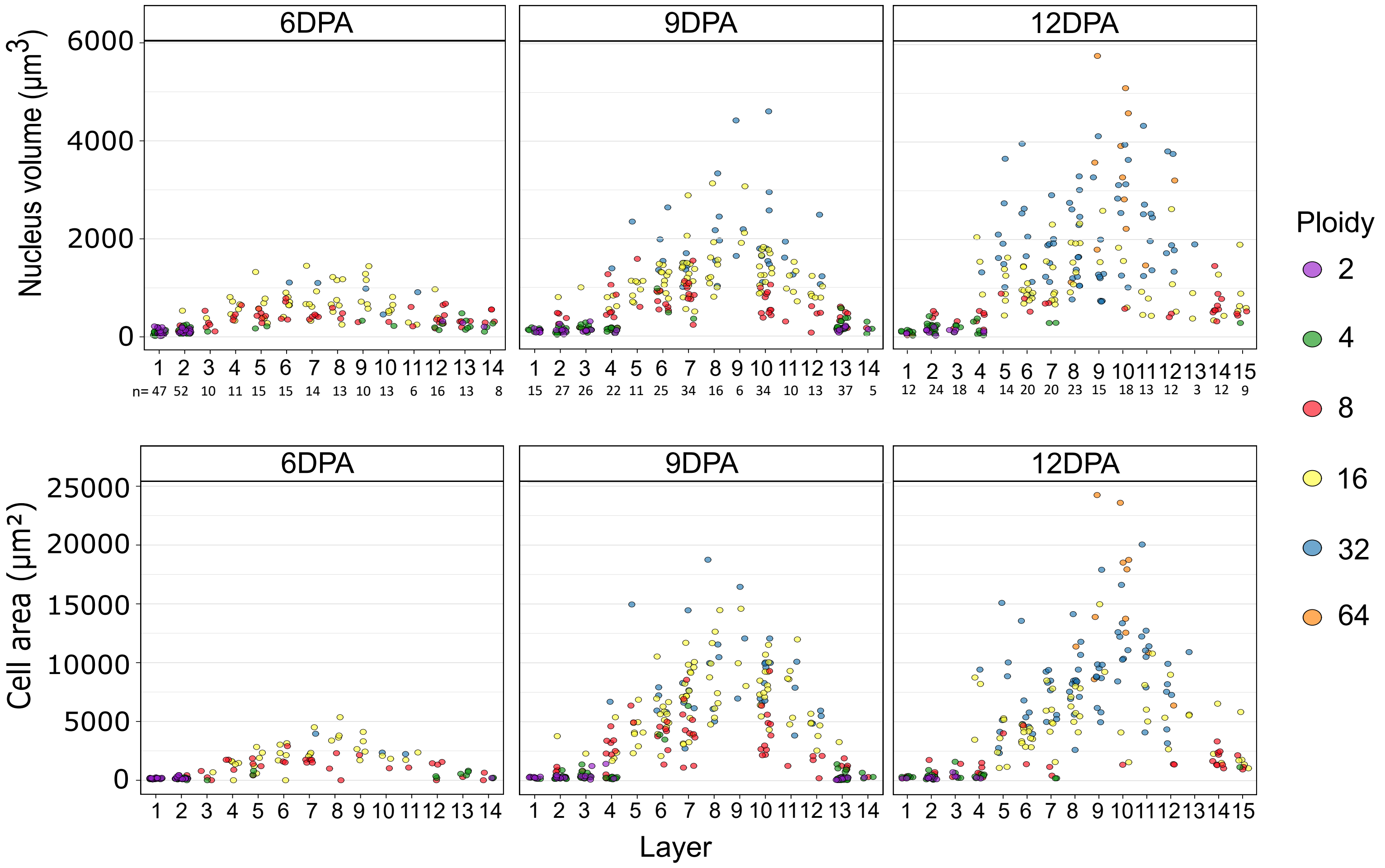


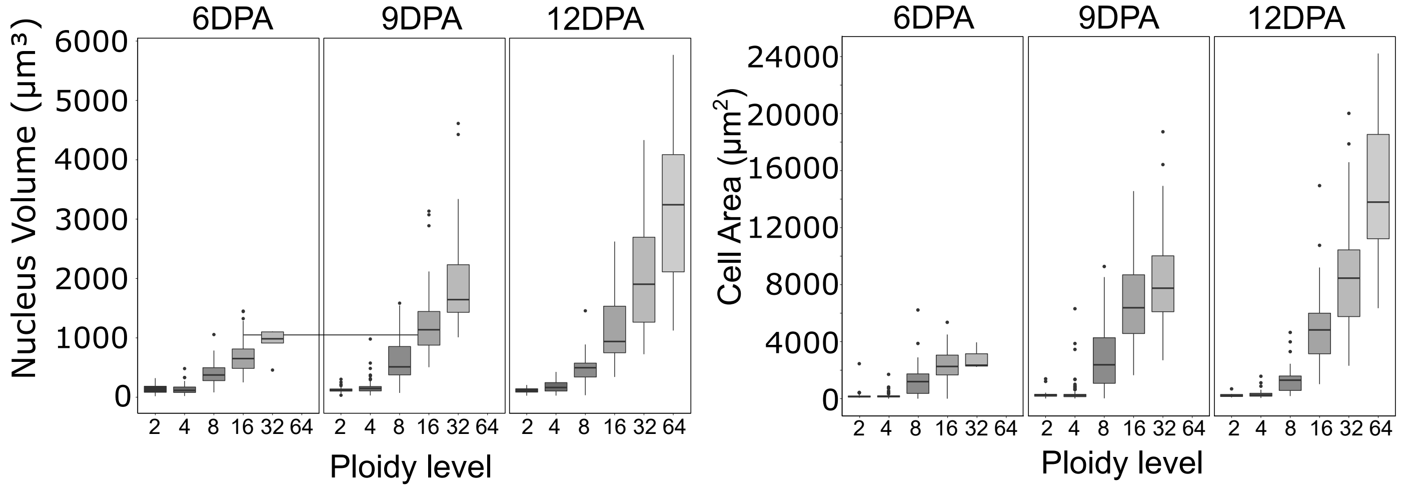

### Figure S7

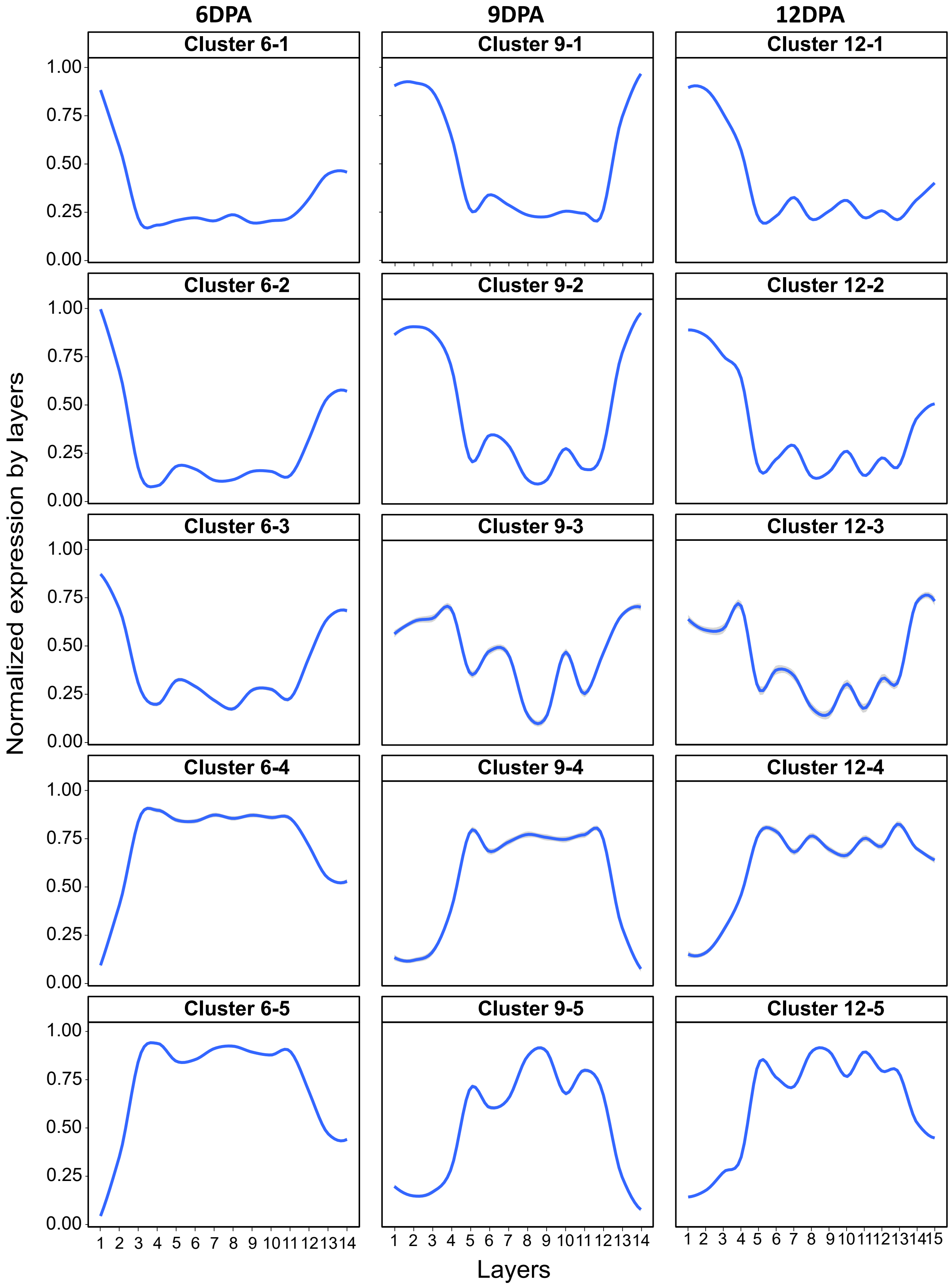
